# Deciphering the links between metabolism and health by building tailored knowledge graphs: application to endometriosis and persistent pollutants

**DOI:** 10.64898/2026.03.02.709027

**Authors:** Meije Mathé, Guillaume Laisney, Olivier Filangi, Franck Giacomoni, Maxime Delmas, German Cano-Sancho, Fabien Jourdan, Clément Frainay

## Abstract

**Motivation:** Knowledge graphs (KGs) are a robust formalism for structuring biomedical knowledge, but large-scale KGs often require complex queries, are difficult for non-experts to explore, and lack real-world context (such as experimental data, clinical conditions, patients symptoms). This limits their usability for addressing specific research questions.

**Results:** We present Kg4j, a computational framework built on FORVM (a large-scale KG containing 82 million compound-biological concept associations), that constructs local, keyword-based sub-graphs tailored to address biomedical research questions. Resulting graphs support hypothetical relationships and can integrate experimental datasets, enabling the discovery of plausible but yet unknown connections. Starting from a conceptual definition of a research field of interest (e.g., disease, symptoms, exposure), the framework extracts relevant associations from FORVM and identifies potential biological mechanisms and chemical compounds.

We applied this approach to endometriosis, exploring links between exposure to Persistent Organic Pollutants (POPs) and disease risk. We propose a novel validation strategy comparing the resulting sub-graph (2,706 nodes and 23,243 edges, 0.002% of FORVM) with recent scientific literature, showing consistency with known findings while also revealing new hypothetical associations requiring further investigation.We also showed that removing duplicated nodes and edges from the KG improves the proportion of validated nodes (from 8.4% to 16%), doubles the precision (from 0.085 to 0.197) while maintaining the recall (0.954 to 0.952), illustrating a trade-off between the loss of potentially relevant but redundant information and the reliability of remaining associations.

By combining automated knowledge mining with experimental data integration, this framework supports reproducible, context-based exploration of biomedical knowledge and systematic hypothesis generation. Applied to endometriosis, it highlights potential mechanisms linking exposure to POPs to the aetiology of the disease, offering a scalable strategy for constructing disease-specific KGs.

**Availability:** The code and data underlying this article are available in the MetExplore repository at https://forge.inrae.fr/metexplore/kg4j.

**Key Messages:**

- Kg4 builds targeted knowledge maps from large biomedical databases using simple keywords.
- Keyword-driven exploration reveals the most relevant disease–exposure relationships without navigating millions of connections.
- Applied to endometriosis, the method recovered known links with persistent organic pollutant exposure.
- Removing redundant information and formatting Knowledge graph as Labeled Property Graph improves the reliability of extracted knowledge.

## Introduction

### Knowledge Graphs in biomedical research

Biomedical research generates massive amounts of data and knowledge from various sources, including scientific literature, clinical records, and omics studies. The amount of information involved requires automated collection, harmonization, and modeling of knowledge to make it usable. In addition to this technical challenge, exploiting this data also faces the cognitive limits of large-scale reasoning.

Traditional relational databases have been used to store and manage biomedical data, but they rely on rigid data schemas that often fail to capture the interconnection and dynamic nature of biological data (7). KGs offer an alternative solution, allowing for the seamless integration of diverse data sources and preserving relationships between entities. They not only facilitate data integration but also support high-level reasoning and inference, and enable the explicit representation of complex relationships and semantic context, making them an essential tool for biomedical research (6) HYPERLINK \l "bookmark32" (26). KGs are used to integrate, manage, and extract large-scale data from different sources and in various formats. They propose a representation of knowledge modeled as a graph. A KG can be represented as a set of subject-predicate-object triples, where each (*s, p, o*) triple represents a relationship *p* between the subject *s* and the object *o*.

Recent work has demonstrated the importance of disease-specific KGs in various medical domains. In 2021, Huang and collaborators presented the construction and use of a KG specific to Kawasaki disease (19), a vasculitis syndrome affecting children. Their KG integrates various knowledge resources, such as clinical guidelines, clinical trials, drug knowledge bases, and medical literature, to support clinical decision-making and research on Kawasaki disease. The constructed KG provides a comprehensive and structured data foundation that enhances the efficiency of literature retrieval and exploration of various aspects of the disease. The authors demonstrated that the integration of various sources of knowledge into a unified graph facilitates more effective clinical decision support, helping researchers and clinicians to quickly access and use relevant information. Similarly, Man and colleagues proposed a disease-specific KG for Kidney Stone Disease (KSD) to enhance visualization, representation and utilization of dispersed knowledge across multiple databases (29). They successfully linked diverse medical data sources to enable efficient semantic search and knowledge discovery for KSD research. More recently, Correira and colleagues (8) constructed a patient-centered application to support epilepsy patients, caregivers and clinicians in self-management and decision making. In this case, the KG is used as a tool to improve access to relevant, individualized information, aiding epilepsy patients and caregivers in managing the condition effectively.

Despite these disease-specific applications, unified frameworks are needed to ensure interoperability and reproducibility across the different disciplines within biomedical research. In 2023, Lobentanzer and collaborators addressed this challenge by introducing a unified framework for data integration and structuring in KG creation. They implemented this standardization in BioCypher (25), a framework designed to transparently build biomedical KGs while keeping the information on the origin of source data. This new development also emphasized the importance of mapping knowledge onto biomedical ontologies to balance human and machine readability and accessibility for non-specialist researchers. Complementing the BioCypher framework, Dreo and colleagues developed OntoWeaver (12), facilitating the reproducible mapping of tabular data into semantic KGs. This tool supports various database formats and enhances the fusion of large-scale information, particularly in the biomedical field.

Regarding the technological foundation of biomedical knowledge graphs, the Semantic Web framework has been widely adopted by major data providers in biomedical research, leading to a large web of life-science linked open data. Researchers can query large federations of decentralized knowledge resources to extract relevant information to contextualize results. We have contributed to this web of knowledge through the FORVM knowledge graph (11), a literature-based resource that links chemicals and biomedical concepts to aid in the interpretation of complex metabolic signatures. It integrates several RDF datasets and leverages the capabilities of the Semantic Web framework to apply ontological reasoning, inferring new relationships between entities and providing diverse levels of abstraction with the potential for novel hypotheses.

Despite the relevance of many large-scale knowledge graphs, some limitations arise in specific applications:

The size of massive knowledge graphs can impact the performance of information retrieval and data mining approaches. Moreover, the infrastructure required to host such resources is costly, often discouraging the deployment of private local instances, which makes the system a one-way, top-down approach with centralized knowledge curation, often lacking user contribution, and with unidirectional data flow. Overcoming this limitation would allow the inclusion of hypotheses and unpublished data, including sensitive or private clinical data, in the analytical pipeline. In addition to challenges with automated processing, human interaction with knowledge graphs is also difficult, from the steep learning curve of the SPARQL query language to the complex visualization of triple-based data, which often results in visually complex and counterintuitive graph structures.

Recently, the community has leaned toward Labeled Property Graphs (LPGs), particularly using the Neo4j system ^1^. Although a full comparison of the advantages and disadvantages of each system is beyond the scope of this article, the Neo4j LPG approach helps to address some of the limitations mentioned above. First and foremost, by allowing links to bear attributes, the LPG tend to be of reduced size compared to their triple-based counterparts, easing visualization and navigation. However, solutions based on LPG have been primarily applied in a bottom-up manner by aggregating in-house data and small- to medium-scale flat datasets, notably using the BioCypher suite. This approach often overlooks valuable information available from the Semantic Web. Although specialized KGs have shown promising results, their construction through the extraction of large-scale knowledge graphs or federations remains challenging. We propose Kg4j ^2^, a Java library aiming to bridge these two approaches by streamlining the extraction and construction of study-scale knowledge graphs from the linked open data supported by the Semantic Web stack, and converting them into BioCypher-compatible LPGs.

### Proof of concept : exposure to POPs and risk of endometriosis

Endometriosis is a chronic inflammatory gynecological disease affecting 5-15% of individuals assigned female at birth, predominantly those of reproductive age (16). It is characterized by ectopic endometrium gowth in the pelvic or abdominal cavity, causing pelvic pain, infertility, dysmenorrhea (painful menstruation), dyspareunia (painful intercourse), dysuria (painful urination), and dyschezia (painful bowel movements) (32), significantly reducing quality of life of affected individuals (22), and increasing the risk of other chronic diseases (21).

The gold standard for diagnosing endometriosis is laparoscopy, but it may not detect deep lesions hidden by adhesions or in the sub-peritoneal space (10). Treatments for endometriosis include surgical intervention and medical therapies such as luteinizing hormone-releasing antagonists or Danazol, although symptoms reappear after treatment in 75% of cases (16).

Recently, approaches using AI have been developed to find new biomarkers of endometriosis, assist with Magnetic Resonance Imaging and Trans-vaginal Ultrasound image analysis, and manage patient reports (13), using omics, imaging, or textual data from patient records. Cetera and colleagues examined how AI could enhance clinical practice (improving diagnosis, treatment planning, and patient management) and discussed potential benefits and limitations (5).

Although physiological mechanisms provide crucial insights into endometriosis pathogenesis, emerging evidence suggests a role for POPs in the onset and progression of the disease. In 2019, Cano-Sancho and collaborators published the first systematic review ever conducted on the relationship between exposure to POPs and the risk of endometriosis in epidemiological studies (3). This meta-analysis revealed positive associations between exposure to organochlorine chemicals - particularly dioxins, Polychlorinated Biphenyls (PCBs), and pesticides - and the risk of endometriosis. Kahn and colleagues hypothesized that prenatal exposure to Endocrine Disrupting Chemicals (EDCs) could lead to reproductive conditions in women, including endometriosis, and suggested associations between Per and poly-fluoroalkyl Substances (PFAS) and endometriosis (20). More recently, Matta and collaborators identified several pollutants associated with endometriosis, including octachlorodibenzofuran (OCDF), heptachlor epoxide, and 3,3’,4,4’-tetrachlorobiphenyl (PCB77) (31). The same authors also provided a systematic review of experimental evidence supporting that tetrachlorodibenzodioxins (TCDDs) promote the onset of endometriosis, lesion growth, cell migration and invasion, and play a role in mediating the steroidogenic disruption and the inflammatory process (30).

Recent metabolomic evidence suggests that certain persistent organochlorine chemicals may contribute to endometriosis through metabolic disruption, linking exposure to trans-nonachlor with altered levels of 2-hydroxybutyric acid in women with deep endometriosis (24).

### Knowledge Graph for endometriosis research

So far, there is little to no unified mechanistic understanding of how specific POPs may contribute to the disease onset or progression. Research on POPs spans diverse research areas, from toxicology to molecular biology and epidemiology. Information about their mechanisms of action, biological targets, and health outcomes is scattered across thousands of publications, and summarizing this dispersed knowledge manually is time consuming. Literature mining, the automated extraction of structured knowledge from scientific papers, provides a way to extract and organize relevant information from a vast source of knowledge (39). Such approaches could help uncover implicit relationships relevant to POPs modes of action, across distinct fields of research.

We propose a KG approach to integrate heterogeneous data sources, enabling the extraction, prioritization, and filtering of relevant information (relations, paths, subgraphs, etc.). More broadly, we developed a command-line tool to construct keywords-based biomedical KGs focused on specific diseases, conditions, or symptoms. Results from experimental metabolomics assays can also be integrated in the KG, to characterize the metabolomic signature for a given condition, and to inform on potential modes of action of chemical stressors.

Endometriosis is an ideal use case for the application of KG-based approaches: this condition remains relatively underexplored compared to other chronic diseases, making the scale of available data suitable for a first Proof Of Concept study. KG approaches in endometriosis and POPs research could help identify novel biomarkers and non-invasive diagnostic tools, while also generating hypotheses regarding the mechanisms underlying disease onset and progression. In addition, recent experimental data from metabolomics studies (31) can be incorporated into the KG, enabling the integration of molecular evidence into the graph. Our method is suited to characterizing associations between POPs exposure and the risk of endometriosis, providing a structured knowledge base for hypothesis generation and mechanism exploration. We used our tool to successfully build a network focused on endometriosis, which we then analyzed to assess the key chemical compounds identified in the KG and compared these findings against existing literature to validate our method.

In the following sections, we first outline the methodology for the KG construction, including the data integration process. Next, we present a proof of concept on endometriosis and POPs, illustrating the usability of this approach to support more effective biomedical research and to assist decision-making. Lastly, we describe the protocol for the validation and analysis of the endometriosis-POPs network, identifying relevant information and its alignment with existing literature.

## Materials and Methods

### RDF data to application specific LPG

The first challenge we address is the transition from heterogeneous RDF datasets to an application-specific biomedical KG as a LPG (Figure 1). The RDF format organizes data into triples, which aligns well with the KG structure. While RDF provides the basis for semantic data representation, RDF databases from multiple sources often have inconsistent modeling (variations in how entities and relationships are represented across sources), and the lack of flexibility of the format (*e*.*g*., blank nodes or reification) can make graphs more complex and harder to navigate. Raw RDF graphs are also impractical to visualize, and for larger graphs, performance and storage become major challenges compared to KG solutions (27) (14).

**Figure 1.**
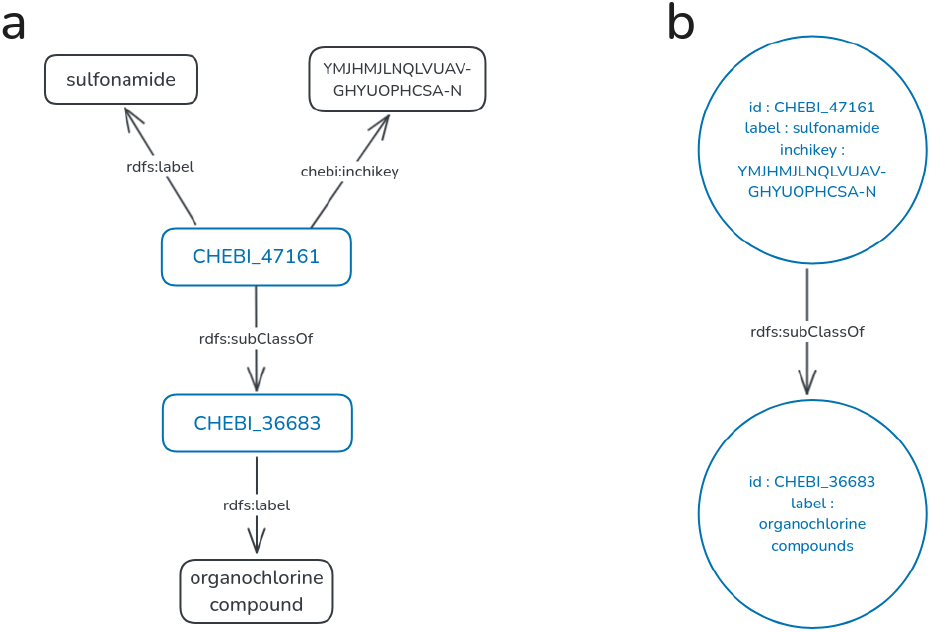
Illustration of the transformation from an RDF representation to a LPG. **(a)** RDF graph describing the sulfonamide compound and **(b)** its corresponding LPG representation.

Reorganizing RDF graphs into application-specific KGs (*e*.*g*., by adding labels or grouping nodes) allows better integration of heterogeneous data, with a flexible schema and semantic richness capturing the meaning of the data. KGs allow automatic inference of new facts, are readable by humans and machines, and can be linked to experimental data. Beyond just storing data, they also organize and contextualize it.

### Kg4j library

We developed Kg4j, an open-source Java library (version 17.0.18) dedicated to the construction, edition, and analysis of KGs focused on biomedical data^3^. This library extends the functionality of the Met4j^4^ library, which is designed for the structural analysis of metabolic networks, to KG objects. We also used the JAVA framework Jena (version 4.10.0) (2), to build semantic web and linked data applications, and the JGraphT library (version 1.5.2) (33) to manage graph theory data structures and algorithms.

The library exports KGs in JSON files, that can then be processed by Neo4j to visualize and analyze the graph.

### Data sources

#### FORVM

The FORVM KG (11) integrates data from various databases, including PubChem RDF compound graphs (chemical compounds), PubMed literature representations (scientific publications), and ontologies such as ChEBI (dictionary of molecular entities) (28), ChemOnt (functions and actions of chemicals) ^5^, CiTO and FaBiO (bibliographic resources and citations) (36), Cheminf (chemical structure and properties) (18), SIO (descriptions of scientific entities) ^6^, and SKOS (any type of controlled vocabulary) (34), as well as the MeSH thesaurus (NLM controlled vocabulary thesaurus used for indexing articles for PubMed) ^7^ representation. These sources provide chemical properties, hierarchies, bibliographic resources, and relationships between compounds and articles. An RDF graph (PMID-CID), established using NCBI-Eutils Elink utility, defines the connections between PubChem compounds and PubMed articles and facilitates the extraction of compound-to-concept links.

FORVM is integrated into a triple-store and accessed via a SPARQL endpoint, enabling complex queries, exploiting ontology-defined semantics, and ensuring data accessibility based on FAIR (Findable, Accessible, Interoperable, and Reusable) principles. This endpoint presents data in the RDF format, a directed, labeled data format for representing information on the web.

#### FORVM dataset extraction

To extract information from the FORVM KG, and create a smaller, local KG focused on chemical compounds and biomedical concepts associated with endometriosis, we used SPARQL. SPARQL is the standard query language and protocol for Linked Open Data on the web or for RDF triplestores, is highly flexible and can query complex relational structures.

To create the KG linking biomedical concepts and chemical compounds, we assembled SPARQL queries to retrieve information from FORVM, related to MeSH descriptors and compounds from the PubChem and ChEBI datasources. The queries’ workflow is described in Figure 2.

**Figure 2.**
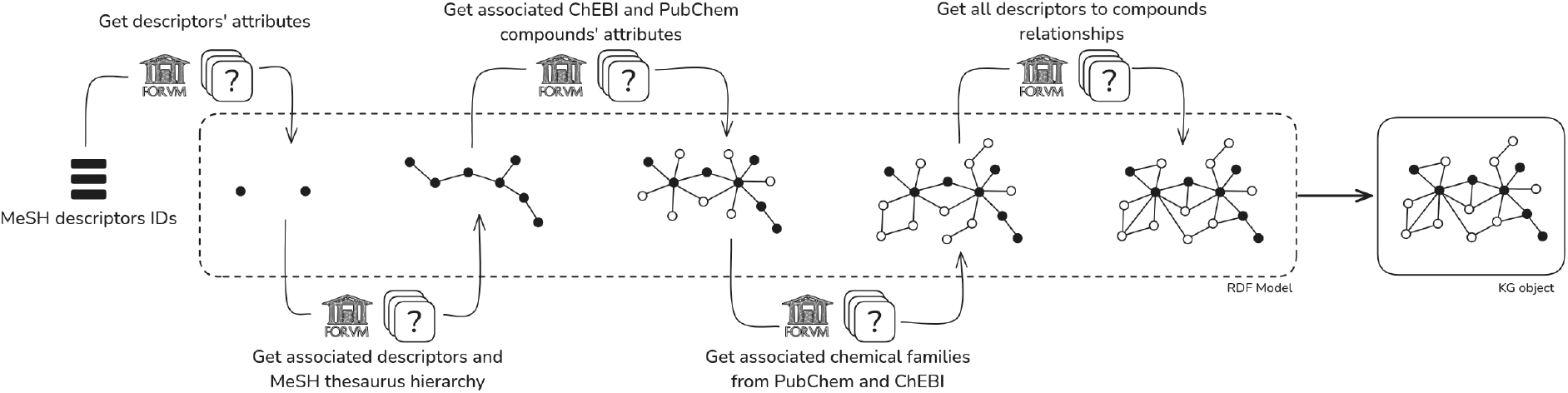
Workflow of SPARQL queries used to build the graph from FORVM. Starting from a list of MeSH descriptors identifiers, descriptors’ attributes (label, id, and active status) are first retrieved and stored in an RDF model. The model is then expanded with associated descriptors and hierarchical relationships derived from the MeSH thesaurus. Next, chemical compounds (from ChEBI and PubChem) associated with the entry descriptors, and their attributes (id, label, source) are added to the model. The chemical families of this compounds are then also retrieved. Lastly, all descriptor-compound relationships are retrieved to ensure completeness of the model. The resulting RDF model is then converted into a KG object represented as a LPG.

### Validation dataset

To establish a validation baseline for the constructed graph, we manually extracted 105 biomedical concepts and 134 chemical compounds (239 entities) from a comprehensive literature review (30). From the article’s findings, we identified relevant entities not initially present in our KG. These were added as artificial nodes to enrich the validation dataset and have a complete representation of the expected knowledge. We then compared the contents of our KG to the validation dataset and used Fisher’s exact test to confirm the significance of the enrichment in the knowledge discussed in the literature, among the entities captured by our KG.

## Results

### Development of the Kg4j Java library for the construction of biomedical KGs

#### KG construction

We developed classes and methods to query FORVM, and to build and export the constructed KG using Java. The library provides several functionalities, including: (1) loading RDF data from a file or an RDF endpoint into a Jena RDF model; (2) executing a combination of SPARQL queries to retrieve specific information (descriptors and compounds labels, names, and hierarchical relationships between them) from the RDF data in the FORVM triplestore; (3) constructing a graph based on the query results using custom builders; (4) exporting the constructed graph to a JSON file or importing a graph from a JSON file; and (5) performing various operations and analyses on the graph using JGraphT algorithms.

The tool allows users to create custom graphs by combining multiple SPARQL queries, from a set of MeSH descriptors semantically describing the research question. It also supports both union and intersection modes, to construct graphs reflecting either the combined or shared queries’ results from the selected descriptors. We defined two main types of node objects in the JGraphT graph object, BioMetaboliteNode and BioConceptNode, representing chemical compounds and biomedical concepts respectively.

Details about the formats used can be found in supplementary materials.

### Graph visualization and analysis

We used Neo4j (1), a graph database platform to store, manage and query the KGs we constructed. Neo4j leverages the Cypher language, a declarative syntax tailored for graph operations, enabling efficient matching and graph manipulation. We added the GDS library ^8^ and the APOC library^9^. The GDS plugin proposes advanced algorithms for graph analysis (community detection, path finding, centrality measures …). The APOC plugin extends Cypher with utility procedures and functions for complex data transformation and integration tasks. We provide a panel of Cypher queries to construct, curate and analyze biomedical KGs in Neo4j. These queries can be found on the GitHub of the Kg4j project, and support data import from a JSON file, semantic node classification, relationship enrichment, and some queries for graph pruning and sparsification. We also included in these queries analytical workflows for structural exploration, computing of centrality and clustering metrics, community detection, and subgraph extraction. Overall this panel enables the exploration of complex networks and the addition of new hypothetical relationships in the KG.

### Validation use case : KG focused on endometriosis and organochlorine compounds

#### KG construction

To demonstrate the applicability of our approach, we present a use case illustrating the construction of a KG based on the endometriosis and OCCs keywords. To build a reference for validation, we manually extracted biomedical concepts and chemical entities discussed in the comprehensive review from Matta et al. (30). This review investigates associations between OCCs exposures and endometriosis.

We then built a KG using Kg4j, based on two entry keywords : Endometriosis (MeSH ID *D004715*) and Hydrocarbons, chlorinated (MeSH ID *D006843*). From the review’s findings, we identified 239 entities. 11 of these entities (5.5%) were not initially present in our endo-OCCs KG and were added as artificial nodes to correctly represent the expected knowledge. Each node was assigned an identifier (MeSH ID, PubChem ID, or ChEBI ID) if available, and a *validated* property to track its source: “*graph*” for all nodes present in the constructed KG; “*true*” for nodes present in the constructed KG and supported by the literature review; and “*false*” for artificial nodes discussed in the review but not found in the KG.

A schematic overview of the built KG’s contents is showed in figure 4.

The endo-OCCs KG contains 2,706 nodes (308 biomedical concepts and 2,398 chemical compounds) and 23,243 edges and is represented in figure 5.

#### KG validation

Artificial nodes were connected both to anchor descriptors (Endometriosis and OCCs) using *skos:related* relationships, and their broader context was added if applicable through SPARQL queries on the FORVM triplestore (e.g. *meshv:broaderDescriptor* relationships).

Additionally, we extended the validation set with mappings between PubChem and ChEBI by integrating broader classes (*rdfs:subClassOf* relationships) and equivalent compounds (*rdf:type* relationships). We also propagated the validation to MeSH narrower descriptors when applicable.

There are two main hubs in the endo-OCCs KG. The first one groups most nodes associated with endometriosis, along with nodes shared with OCCs and the second hub groups nodes associated with OCCs. The first hub is characterized by a higher density (density values are depicted in figure 8), in part due to the hierarchical structure of the MeSH thesaurus.

Out of the 2,706 nodes in the KG, 554 (20.5%) are directly related to endometriosis, while 1,285 nodes (47.5%) are directly related to OCCs. 416 nodes (15.4%) are common to both the endometriosis and OCCs ecosystems. The remaining 867 are indirectly connected to either entry concepts, and often include broader entities such as chemical classes or higher level MeSH descriptors.

The nodes validated through the literature review are mainly found in the most interconnected part of the KG (139 out of 228 - 61% - in the “endometriosis” ecosystem of the graph), suggesting they represent well-established entities that are frequently co-mentioned or thoroughly described in scientific literature. The non-validated nodes tend to represent more inferred associations, including artificial nodes. In total, 228 of the 2,706 nodes (8.4%) were discussed in the literature review, and 228 out of the 239 entities (95.4%) from the review are present in the KG.

#### Graph pruning

The presence of duplicated nodes and edges, and the high density of the KG can introduce topological bias in downstream analyses and complicate its exploration and visualization. We applied a process of graph pruning, consisting of removing unnecessary nodes or edges while maintaining functionality in the graph.

The pruning steps we applied on the KG are described in figure 6. Notably, we removed entry nodes (input MeSH descriptors) : they are linked to almost all the other nodes in the graph, and tend to skew the topology of the graph by artificially increasing centrality and make it harder to observe the natural structure of edges. We also removed duplicated nodes and edges.

We removed isolated nodes at each step and monitored graph metrics such as diameter, density and clustering coefficient across the pruning steps. The diameter measures the longest shortest path between any pair of nodes in the graph, reflecting its overall size and connectivity. The density captures how close the graph is to being fully connected. The clustering coefficient measures how much nodes tend to cluster together, revealing the presence of communities. Monitoring these metrics allows us to assess how pruning impacts the global and local patterns in the graph. The Cypher queries we used to prune the graph and compute graph metrics are in the queries library.

The graph pruning process led to a significant reduction in both node and edge counts (Figure 7 a and b). Pruning initially increased the graph’s diameter and average path length, due to its fragmentation, followed by a slight increase and stabilization in the later steps. The loss of triangles and higher-order cliques (Figure 7 d) confirms the reduction of local density and community structures. The relationships *rdf:type* (representing identical comopounds between PubChem and ChEBI) disappear from the KG after the first pruning step, which is expected. The whole graph and the final pruned graph are presented in figure 8.

The pruned KG contains 1117 nodes - 305 biomedical concepts (27.3%) and 812 chemical compounds (72.7%) – and 7,849 edges. Among these nodes, 431 (38.6%) were directly linked to the endometriosis concept, 529 (431%) were directly linked to the OCCs concept, and 235 (21%) were shared between both ecosystems. This reduction mainly affected weakly connected nodes : out of the 1589 discarded nodes, 998 (62.8%) were removed because they became isolated after a pruning step, and not because of the intended pruning process itself. A greater proportion of nodes (33.3%) from the OCCs ecosystem than the endometriosis ecosystem (4.5%) were excluded during the pruning process due to their limited connectivity: they were typically associated with only one biomedical concept, representing OCCs. Of all nodes, 179 (16%) were discussed in the literature review, with only one artificial node remaining, the biomedical concept *Disease Progression*. The proportion of validated nodes thus doubles in the graph after the pruning process, illustrating a trade-off between the exhaustibility and the quality of retrieved information.

#### Validated node enrichment

We considered all entities discussed in the literature review as the set of validated nodes. We performed Fisher’s exact tests (42) to assess whether the validated nodes were significantly overrepresented in the endo-OCCs KG, comparing the numbers of validated nodes represented in the graph or missed during the KG building process. We performed these tests on both the whole and pruned graphs.

The overlap between our graph and the validation set is represented in a Venn diagram, before and after pruning (Figure 9), highlighting the intersection of previously published knowledge with our graph.

We found a significant enrichment of validated nodes in both graphs relative to the overall validation set (Fisher’s exact test, *p − value <* 0.05). However, due to the extremely large size of the background universe (all entities in FORVM), the Fisher’s test p-value alone provides limited interpretability, as even small overlaps could yield statistical relevance.

We computed additional complementary metrics to obtain a more informative overview of the overlap: **fold enrichment**, quantifying how much greater the observed overlap is compared to a random overlap; **Odds Ratio** (40), assessing whether membership in the KG increases the likelihood of being mentioned in the review; **precision**, the proportion of KG’s entities that are mentioned in the review; **recall**, the proportion of entities mentioned in the review that belong to the KG (37). The metrics’ values are displayed on Figure 9.

Overall, the fold enrichment, the Odds Ratio, and the precision increase after pruning, while the recall is maintained. This suggests that pruning reduces noise and increases signal. The graph becomes smaller but more informative, and is enriched for validated entities without losing a significant amount of coverage.

#### Hubs

We identified high-centrality hubs (highly connected nodes) in the whole endo-OCCs KG, representing the semantic landscape of research on OCCs and endometriosis. The central chemical entities were steroids (*3-oxo-Delta(4) steroid, 20-oxo steroid, 17b-hydroxy steroid*,…) and pollutants classes (*PCB and related compounds, PCB, nitrile, chlorobenzene*,…), reflecting existing literature on environmental exposures (31) (30). For biomedical concepts, main nodes were associated with hormones and hormonal therapy (*Hormones, Hormone Substitutes, and Hormone Antagonists, Physiological Effects of Drugs*, …) and physio-pathological processes relevant to endometriosis *(Disease Progression, Pain, Pelvic Pain*, …), highlighting the relation between the disease and potential chemical drivers. The full ranked list of nodes based on their centrality (degree, betweenness and PageRank) are provided in the supplementary data.

The mean degree centrality of the validated nodes is higher than the global average centrality both before (17.18 vs. 60.33) and after pruning (14.05 vs. 24.09). The validated nodes correspond to highly interconnected entities in the graph.

#### Hypothesis generation

While the most central nodes reflect well-established knowledge in the scientific literature, nodes with moderate or lower centrality may represent underexplored or emerging associations and provide valuable insights for hypothesis generation. Such nodes, not particularly important in the topology of the KG, and not discussed in the review, may highlight potential novel mechanistic links.

For example, biomedical concepts including *Cell Transformation, Neoplastic* (degree = 13), *Uterine Cervical Neoplasms* (degree = 22), and *Metaplasia* (degree = 33) suggest a possible link between endometriosis and oncogenic processes, supported in prior studies (23) (38).

Kg4j enables domain expert to refine the KG by incorporating additional evidence, proposing hypothetical relationships between existing entities, or adding new concepts.

## Discussion

We introduced kg4j, an open-source Java framework for constructing and analyzing local custom biomedical KGs. The framework enables the conversion of RDF resources into LPG, improving usability, interpretability and exploration. Through a proof of concept on endometriosis and organochlorine pollutants, we demonstrated how tailored disease-specific KGs can integrate heterogeneous biomedical data to address specific research questions. The framework supports the creation of hypothetical relationships, facilitating the exploration of plausible but unconfirmed biological mechanisms.

This proof of concept demonstrates the importance of knowledge mining in toxicology and reproductive health, where a big part of the information relevant to disease mechanisms remains fragmented in the literature. KGs can uncover potentially important relationships not yet described, expanding the research scope in the study of complex and multi-factorial etiologies, such as endometriosis. The framework could also assist in systematic reviews by alleviating the effort required for the initial screening of the literature, where information is often sparse.

The validation protocol, which combines manual extraction of entities from the literature with statistical enrichment analyses, confirmed that the KG is in line with established scientific knowledge. This shows that the graph captures biologically meaningful associations. The link between pollutant exposure and endometriosis remains an active area of research, which shows the non triviality of our applicative setting. Yet, the generalization of our findings to other areas of research remains to be investigated.

Although not demonstrated in this study, Kg4j is extended to incorporate additional chemical compounds into th KG, such as metabolites identified through metabolomics experiments. Integrating these compounds would enable researchers to characterize condition-specific metabolic signatures and prioritize key metabolites for targeted downstream analysis and further experiments.

We show that graph pruning improved the interpretability of the KG while maintaining essential information. Pruning addressed issues such as redundancy across ontologies (*e*.*g*. PubChem and ChEBI), the over-importance of anchor nodes, and edge duplication, resulting in a more balanced graph.

The integration with Neo4j extended the potential of the framework, enabling complex graph operations such as community detection, clustering, and pruning steps. This makes the KG accessible to non-experts, supporting exploratory analyses. Compatibility with the Biolink model (41) and the KGX ^10^ format makes the graph interoperable with other biomedical KG and tools such as BioCypher or KGHub, broadening its usability.

Despite these contributions, several limitations must be acknowledged. First, the coverage and quality of the KG are limited by the underlying ontologies, which may not contain all relevant concepts or definitions. Second, the validation protocol relied on a single literature review, introducing potential biases in publication selection and coverage. Although FORVM v2021 was used to align with the review’s publication date, some relevant entities may have been missed.

Furthermore, biomedical literature is not a perfect ground truth: some associations may be true but remain unreported, while others may be false. Over-representation of well-studied entities could also inflate their perceived importance in the graph. In addition, interpreting the KG requires domain expertise, especially to distinguish biologically plausible from unfounded associations.

The KG provides a network of high-level biomedical concepts to guide literature-based research, it lacks refined information such as temporal, dose-response, or population-level information, which are critical in toxicology. Moreover, while the KG suggests hypothetical links, these must ultimately be confirmed through experimental or clinical validation.

## Conclusion

This work demonstrates that LPGs provide a more interpretable framework than traditional RDF KGs for biomedical knowledge analysis, enabling richer semantics, flexible queries, and intuitive exploration of complex relationships. Kg4j offers a flexible solution for automated KG extraction and construction, facilitating the integration of heterogeneous biomedical resources with experimental data.

The proof of concept on endometriosis and POPs illustrates this approach : by aggregating diverse data sources, the framework supports knowledge synthesis, hypothesis generation and priorization, and mechanistic insight discovery.

Future developments will enhance the framework with the incorporation of additional data sources to the KG.

Finally, advancing the methodology for graph pruning is essential to balance the complexity and completeness of the graph, ensuring that it remains usable and faithful to the underlying knowledge.

## Abbreviations

KG: Knowledge Graph
AI: Artificial Intelligence
POPs: Persistent Organic Pollutants
RDF: Resource Description Framework
KGX: Knowledge Graph Exchange
GDS: Graph Data Science
APOC: Awesome Procedure On Cypher
OCCs: organochlorine compounds
LPG: Labeled Property Graph

## Data availability

The code and data underlying this article are available at https://forge.inrae.fr/metexplore/kg4j.

## Author contributions statement

MM : formal analysis - investigation - software - wirting (original draft) - data curation. GL, OF and FG : data curation – resources - writing (review). MD : resources - software - writing (review). GCS : validation - writing (review) - funding aquisition, FJ : project administration - supervision - funding aquisition, CF : conceptualisation - data curation - methodology - supervision - project administration - resources - software - writing (original draft + review)

## Acknowledgments

The authors express their gratitude and appreciation to INRAE DIGIT-BIO for their support, Nathalie Poupin and Axel Raux for providing data and valuable insights that supported the evaluation of this approach, and to Juliette Cooke for proofreading the manuscript.

## Supplementary materials

### Usage

#### Interoperability: the Biolink model

The Biolink model (41) is an open-source framework describing how biological entities and their relationships are represented in KGs. It covers entities such as genes, diseases, chemicals, organisms, genomic data and phenotypes to unify biomedical entities and their relationships across datasets. The framework combines object-oriented and graph-based structures to define hierarchical classes representing entities, and predicates representing relationships. It also ensures consistency across databases and proposes guidelines regarding the interaction between entities. For example, in the BioCypher framework (25) and the Biomedical Data Translator (15), the Biolink model is used to characterize the entities and relationships of the constructed KG. We selected the relevant classes and predicates from Biolink to construct our KG, making it interoperable with other KGs built using the BioCypher and KG-Hub frameworks (12; HYPERLINK \l "bookmark12" 4). We selected the following entities and relationships from the BioLink model to describe elements in the graph:

1. ChemicalEntity: represents chemical compounds from the PubChem and ChEBI databases.
2. CommonDataElement: represents MeSH descriptors.
3. BiologicalEntity: represents any other type of node in the graph.
4. broad matches: equivalent to the meshv:broaderDescriptor relationship in MeSH, describes descriptors that have a broader meaning.
5. subclass of: equivalent to the rdfs:subClassOf relationship in ChEBI, and the vocab:has parent relationship in PubChem, describes the class hierarchy between chemical entities (ChEBI and PubChem chemical families).
6. rdf:type: describes the equivalence between chemical compounds defined in both the ChEBI and PubChem databases.
7. occurs together in literature with: equivalent to the skos:related relationship in FORVM, describes a link between two biological entity co-mentioned in the scientific literature.

#### KGX format

To ensure interoperability and fit to established standards (9), the resulting KG is exported in the KGX format ^11^. KGX is a Python libary and set of command line applications designed to exchange KGs aligned with the Biolink model. Exporting the KG in KGX format facilitates integration with other KG frameworks and tools (4) HYPERLINK \l "bookmark36" (35) HYPERLINK \l "bookmark20" (17), supports conversion to multiple formats (RDF, CSV/TSV, Neo4j), and ensures that the graph conforms to Biolink model standars for node categories, edge types, and properties.

#### Command Line Interface

To facilitate the Kg4j library usage, we developed a Command Line Interface. This application constructs a KG based on a set of MeSH descriptors and a list of chemical compounds’ identifiers. The graph is then exported into a JSON file.

The user can define 4 arguments to specify (1) a list of MeSH descriptors describing the field of research; (2) a list of ChEBI or MetaNetX identifiers (optional) representing results from a metabolomics experiment; (3) the KG construction mode (union or intersection) and (4) the name of the output file (Figure 3).

**Figure 3.**
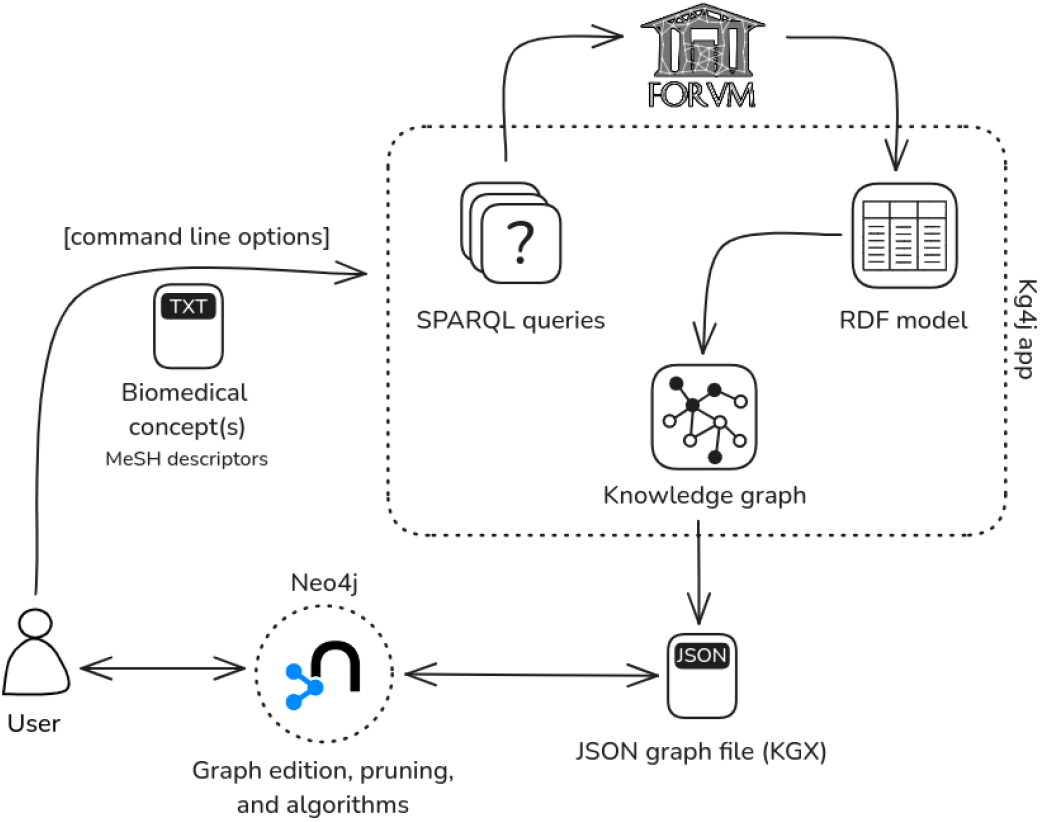
Command-line pipeline for constructing and analyzing the KG. Users provide and input file containing a list of MeSH descriptors identifiers through command line options. The Kg4j application executes SPARQL queries to retrieve relevant information from FORVM, as described in Figure 2. The resulting KG is exported as a JSON file and can be imported into Neo4j for graph visualization, pruning, and analysis.

**Figure 4.**
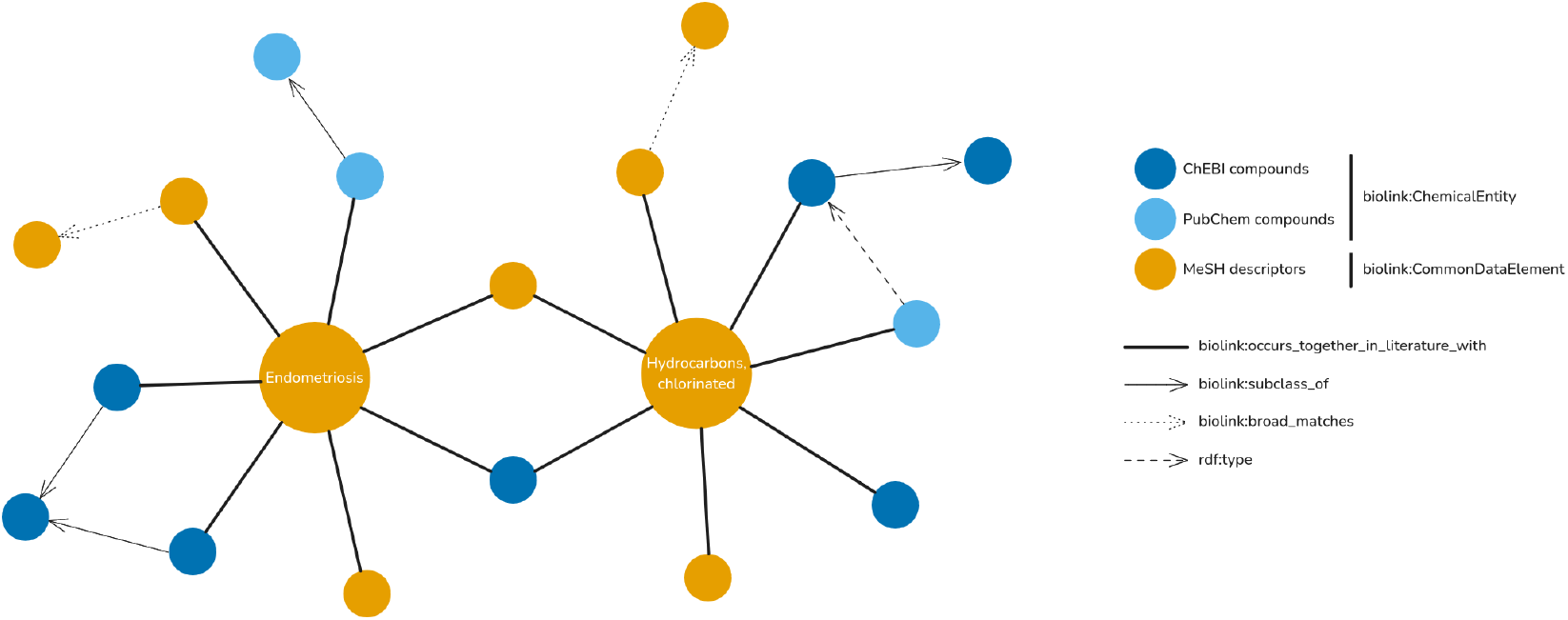
Schematic overview of the knowledge graph structure. Chemical entities from PubChem and PubMed are represented as *biolink:ChemicalEntity* nodes (blue), and MeSH descriptors are represented as *biolink:CommonDataElement* nodes (orange). Edges represent semantic relationships between nodes : *rdf:type, biolink:subclass of, biolink:broad matches*, and *biolink:occurs together in literature with*.

**Figure 5.**
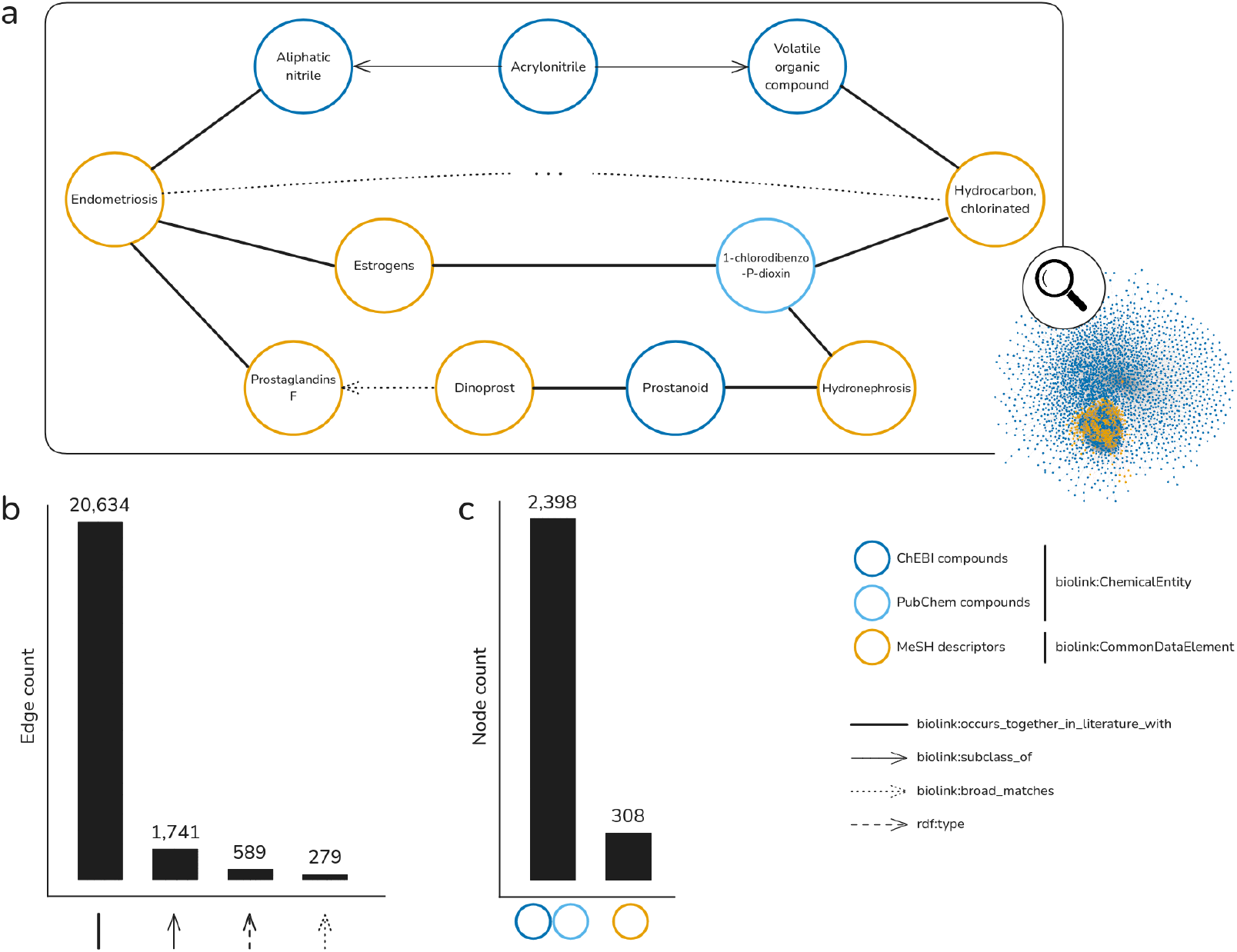
Visualization of a subset of the endometriosis - OCCs KG. **(a)** Subgraph highlighting the different relationships and entities shared between the two core concepts *Endometriosis* and *Hydrocarbons, chlorinated*. Orange nodes represent *CommonDataElement* entities (MeSH descriptors) and blue nodes represent *ChemicalEntity* entities. **(b)** Distribution of edge types across the whole KG (original FORVM). The majority of edges correspond to co-occurrence in literature. **(c)** Distribution of node types in the graph. *ChemicalEntity* (n = 2398) dominate, with a smaller number of *CommonDataElement* (n = 308).

**Figure 6.**
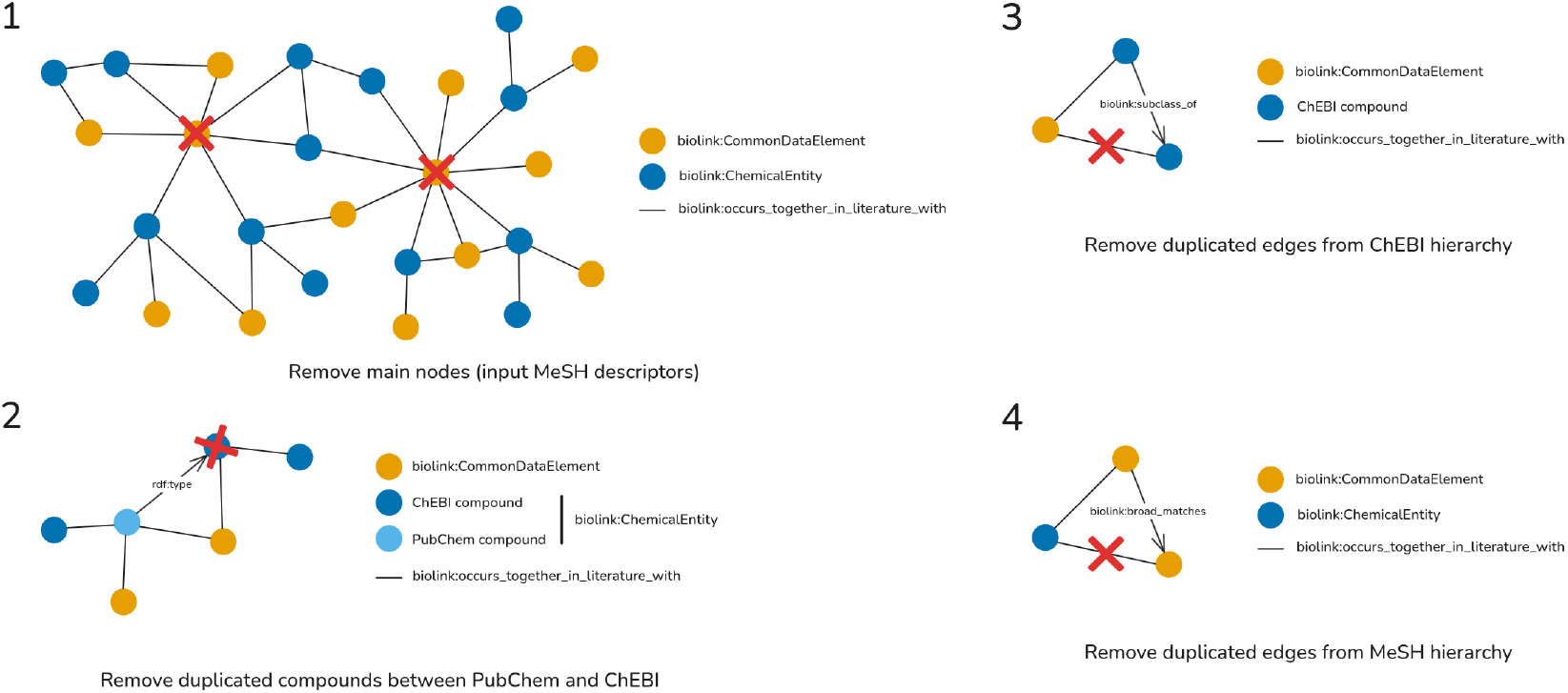
Graphical representation of the different pruning steps applied to the KG.

**Figure 7.**
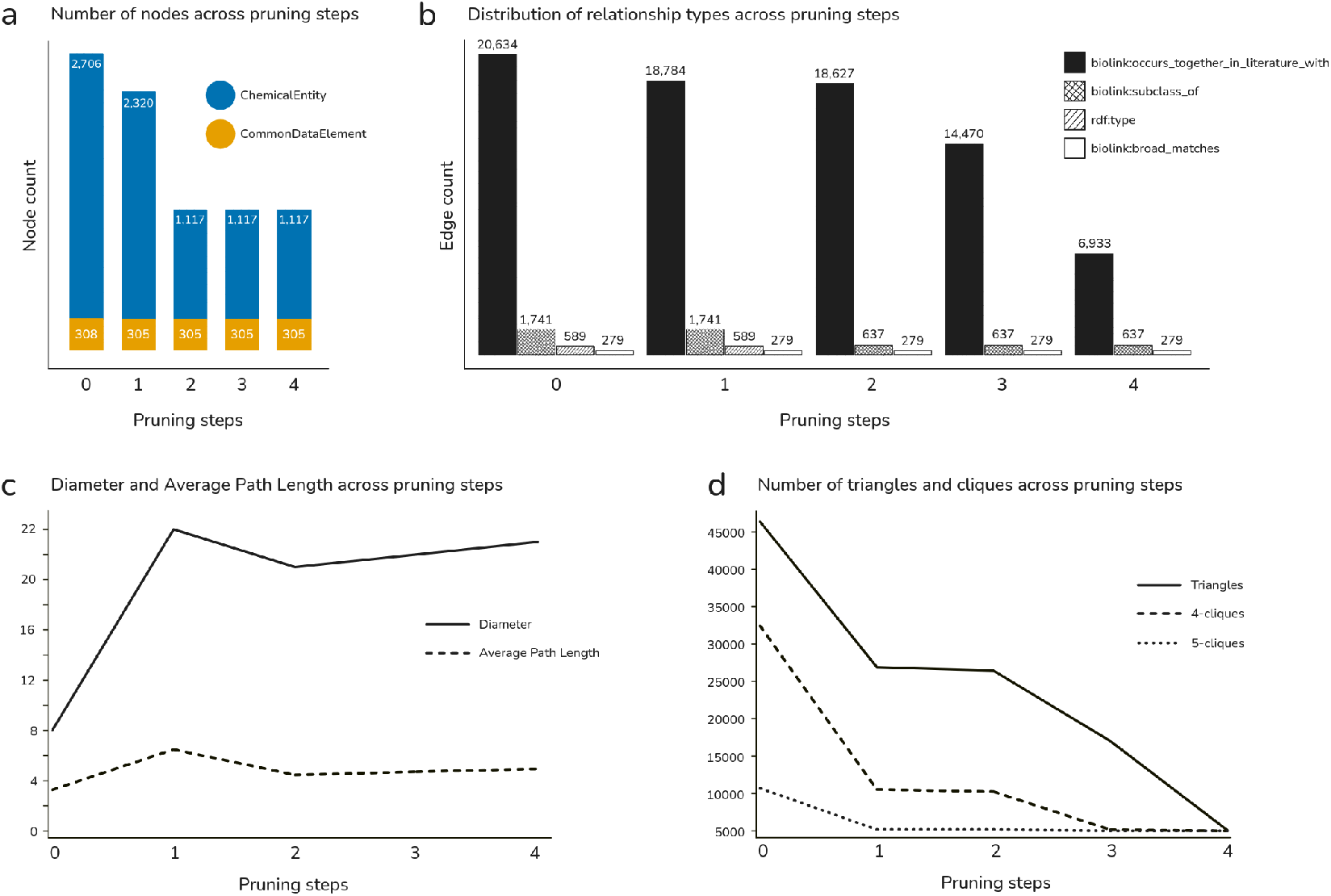
Evolution of graph metrics across pruning steps. **(a)** Evolution of node counts and **(b)** frequency of relationship types across pruning steps. **(c)** Graph diameter and average path length evolution across pruning steps, reflecting connectivity changes. **(d)** Counts of triangles and higher-order cliques (4-cliques and 5-cliques) across pruning, reflecting changes in graph density and local clustering.

**Figure 8.**
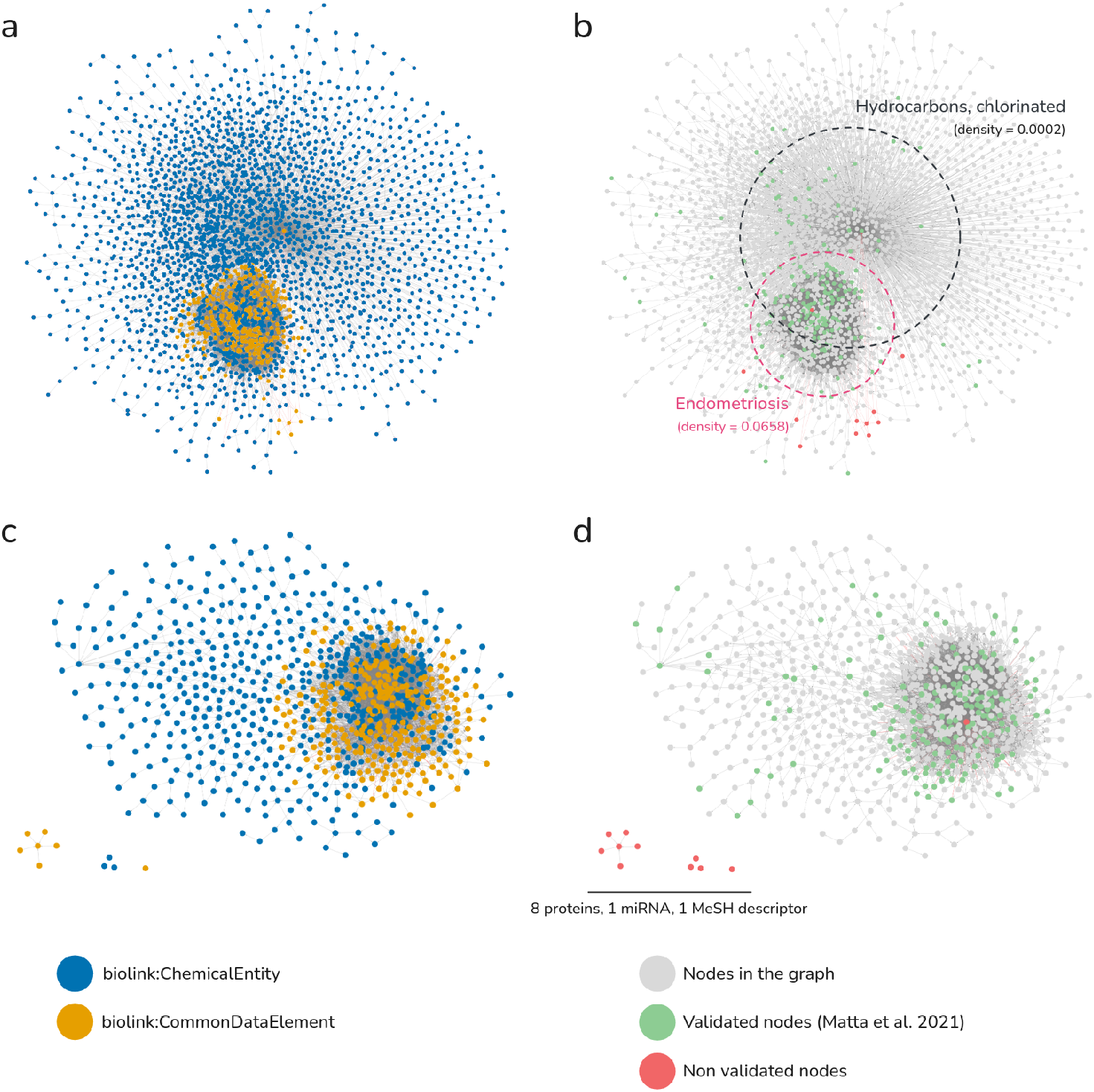
Visualization of the pruned endometriosis - OCCs knowledge with Neo4j Bloom. **(a)** and **(c)** Bipartite graphs illustrating relationships between chemical compounds (blue) and biomedical concepts (orange). **(b)** and **(c)** Representation of the same graphs, highlighting validated nodes (green) corresponding to previously reported knowledge, and non-validated nodes (red). Grey nodes represent all nodes in the graph. The set of isolated nodes at the bottom corresponds to the non validated nodes : 8 proteins, 1 miRNA and a MeSH descriptor (*Incidence*), disconnected from the main graph after pruning. **(a)** and **(b)** illustrate the whole graph, whereas **(c)** and **(d)** illustrate the graph after pruning. Smaller disconnected subgraphs are not represented here.

**Figure 9.**
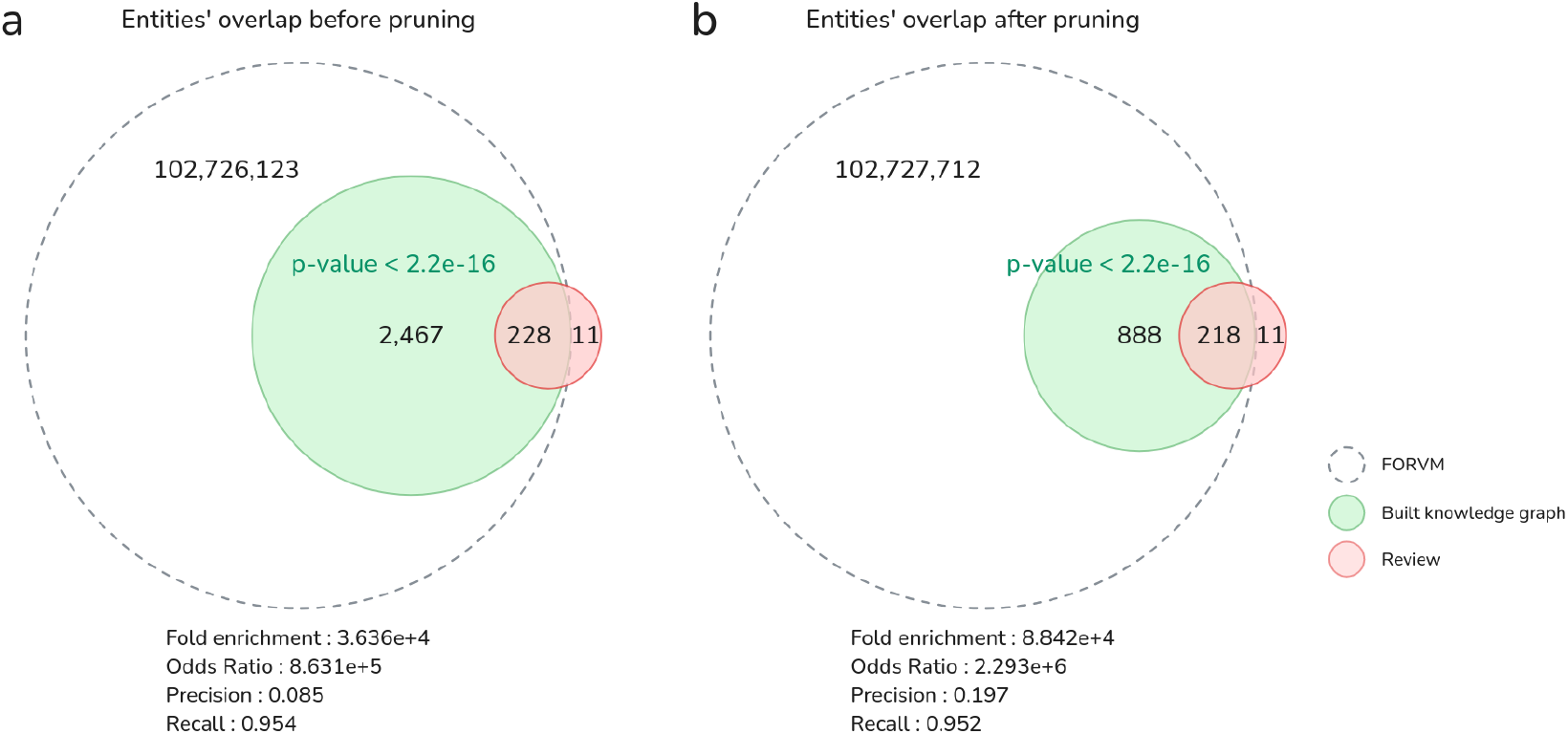
Enrichment of validated nodes in the knowledge graph. Schematic representation of the overlap between entities from the built knowledge graph (green) and the validation set described in the literature review (red), before **(a)** and after **(b)** pruning. The sizes of the circles are not to scale.

## Footnotes

1 https://github.com/neo4j/neo4j

2 https://forge.inrae.fr/metexplore/kg4j

3 https://forge.inrae.fr/metexplore/kg4j

4 https://forge.inrae.fr/metexplore/met4j

5 https://kghub.org/kg-registry/resource/chemont/chemont.html

6 https://github.com/MaastrichtU-IDS/semanticscience

7 https://www.nlm.nih.gov/mesh/meshhome.html

8 https://github.com/neo4j/graph-data-science

9 https://github.com/neo4j-contrib/neo4j-apoc-procedures

10 https://github.com/biolink/kgx

11 https://github.com/biolink/kgx

## References

1. Neo4j documentation - Neo4j Documentation.

2. Apache Software Foundation. Apache jena. https://jena.apache.org/, 2021. Accessed: 2021.

3. German Cano-Sancho, Stéphane Ploteau, Komodo Matta, Evdochia Adoamnei, Germaine Buck Louis, Jaime Mendiola, Emile Darai, Jean Squifflet, Bruno Le Bizec, and Jean-Philippe Antignac. Human epidemiological evidence about the associations between exposure to organochlorine chemicals and endometriosis: Systematic review and meta-analysis. Environment International, 123:209–223, February 2019.

4. J. Harry Caufield, Tim Putman, Kevin Schaper, Deepak R. Unni, Harshad Hegde, Tiffany J. Callahan, Luca Cappelletti, Sierra A. T. Moxon, Vida Ravanmehr, Seth Carbon, Lauren E. Chan, Katherina Cortes, Kent A. Shefchek, Glass Elsarboukh, Jim Balhoff, Tommaso Fontana, Nicolas Matentzoglu, Richard M. Bruskiewich, Anne E. Thessen, Nomi L. Harris, Monica C. Munoz-Torres, Melissa A. Haendel, Peter N. Robinson, Marcin P. Joachimiak, Christopher J. Mungall, and Justin T. Reese. KG-Hub-building and exchanging biological knowledge graphs. Bioinformatics (Oxford, England), 39(7):btad418, July 2023.

5. Giulia Emily Cetera, Alberto Eugenio Tozzi, Valentina Chiappa, Isabella Castiglioni, Camilla Erminia Maria Merli, and Paolo Vercellini. Artificial Intelligence in the Management of Women with Endometriosis and Adenomyosis: Can Machines Ever Be Worse Than Humans? Journal of Clinical Medicine, 13(10):2950, May 2024.

6. Payal Chandak, Kexin Huang, and Marinka Zitnik. Building a knowledge graph to enable precision medicine. Scientific Data, 10(1):67, February 2023.

7. Jake Y. Chen. Biological Database Modeling. Artech House eBooks, October 2007.

8. Rion Brattig Correia, Jordan C. Rozum, Leonard Cross, Jack Felag, Michael Gallant, Ziqi Guo, Bruce W. Herr, Aehong Min, Jon Sanchez-Valle, Deborah Stungis Rocha, Alfonso Valencia, Xuan Wang, Katy Börner, Wendy Miller, and Luis M. Rocha. myAURA: a personalized health library for epilepsy management via knowledge graph sparsification and visualization. Journal of the American Medical Informatics Association: JAMIA, page ocaf012, January 2025.

9. Katherina G Cortes, Shilpa Sundar, Sarah Gehrke, Keenan Manpearl, Junxia Lin, Daniel Robert Korn, Harry Caufield, Kevin Schaper, Justin Reese, Kushal Koirala, Lawrence E Hunter, E. Kathleen Carter, Marcello DeLuca, Arjun Krishnan, Chris Mungall, and Melissa Haendel. Improving Biomedical Knowledge Graph Quality: A Community Approach. ArXiv, page 2508.21774v1, August 2025.

10. Antônio Coutinho, Leonardo Kayat Bittencourt, Cíntia E. Pires, Flávia Junqueira, Cláudio Márcio Amaral de Oliveira Lima, Elisa Coutinho, Marisa A. Domingues, Romeu C. Domingues, and Edson Marchiori. MR Imaging in Deep Pelvic Endometriosis: A Pictorial Essay. RadioGraphics, 31(2):549–567, March 2011. Publisher: Radiological Society of North America.

11. Maxime Delmas, Maxime Delmas, O. Filangi, Olivier Filangi, Paulhe N, Florence Vinson, Duperier C, Garrier W, Saunier P, Y. Pitarch, Yoann Pitarch, Fabien Jourdan, Franck Giacomoni, and Clément Frainay. FORUM: Building a Knowledge Graph from public databases and scientific literature to extract associations between chemicals and diseases. Bioinformatics, September 2021. MAG ID: 3198511414.

12. Johann Dreo, Claire Laudy, Sebastian Lobentanzer, Marko Baric, Ekaterina Gaydukova, and Benno Schwikowski. Reproducible Mapping of Tabular Data into Semantic Knowledge Graphs with OntoWeaver and BioCypher. March 2024.

13. Brie Dungate, Dwayne R Tucker, Emma Goodwin, and Paul J Yong. Assessing the Utility of artificial intelligence in endometriosis: Promises and pitfalls. Women’s Health, 20:17455057241248121, January 2024. Publisher: SAGE Publications Ltd STM.

14. Antonio Fabregat, Florian Korninger, Guilherme Viteri, Konstantinos Sidiropoulos, Pablo Marin-Garcia, Peipei Ping, Guanming Wu, Lincoln Stein, Peter D’Eustachio, and Henning Hermjakob. Reactome graph database: Efficient access to complex pathway data. PLOS Computational Biology, 14(1):e1005968, January 2018.

15. Karamarie Fecho, Gwênlyn Glusman, Sergio E. Baranzini, Chris Bizon, Matthew Brush, William Byrd, Lawrence Chung, Andrew Crouse, Eric Deutsch, Michel Dumontier, Aleksandra Foksinska, Jennifer Hadlock, Kaiwen He, Sui Huang, Robert Hubal, Gregory M. Hyde, Sharat Israni, Kelyne Kenmogne, David Koslicki, Jana Dorfman Marcette, Ewy A. Mathe, Abrar Mesbah, Sierra A. T. Moxon, Christopher J. Mungall, John Osborne, Carrie Pasfield, Guangrong Qin, Stephen A. Ramsey, Justin Reese, Jared C. Roach, Reese Rose, Karthik Soman, Andrew I. Su, Casey Ta, Gaurav Vaidya, Rosina Weber, Qi Wei, Mark Williams, Chunlei Wu, Colleen Xu, and Chase Yakaboski. Announcing the Biomedical Data Translator: Initial Public Release. Clinical and Translational Science, 18(7):e70284, July 2025.

16. Linda C. Giudice, Tomiko T. Oskotsky, Simileoluwa Falako, Jessica Opoku-Anane, and Marina Sirota. Endometriosis in the era of precision medicine and impact on sexual and reproductive health across the lifespan and in diverse populations. The FASEB Journal, 37(9):e23130, September 2023.

17. Skye L. Goetz, Amy K. Glen, and Gwênlyn Glusman. MicrobiomeKG: bridging microbiome research and host health through knowledge graphs. Frontiers in Systems Biology, 5:1544432, 2025.

18. Janna Hastings, Leonid Chepelev, Egon Willighagen, Nico Adams, Christoph Steinbeck, and Michel Dumontier. The Chemical Information Ontology: Provenance and Disambiguation for Chemical Data on the Biological Semantic Web. PLOS ONE, 6(10):e25513, October 2011.

19. Zhisheng Huang, Qing Hu, Mingqun Liao, Cong Miao, Chengyi Wang, and Guanghua Liu. Knowledge Graphs of Kawasaki Disease. Health Information Science and Systems, 9(1):11, February 2021.

20. Linda G Kahn, Claire Philippat, Shoji F Nakayama, Rémy Slama, and Leonardo Trasande. Endocrine-disrupting chemicals: implications for human health. The Lancet Diabetes & Endocrinology, 8(8):703–718, August 2020.

21. Marina Kvaskoff, Fan Mu, Kathryn L. Terry, Holly R. Harris, Elizabeth M. Poole, Leslie Farland, and Stacey A. Missmer. Endometriosis: a high-risk population for major chronic diseases? Human Reproduction Update, 21(4):500–516, July 2015.

22. Claruza Braga Holanda Lavor, Francisca Adriele Vieira Neta, Antonio Brazil Viana, and Francisco das Chagas Medeiros. The impact of surgical treatment for deep endometriosis: metabolic profile, quality of life and psychological aspects. Revista Brasileira de Ginecologia e Obstetrícia, 46:e–rbgo42, June 2024.

23. S. Leenen, M. Hermens, P. J. de Vos van Steenwijk, R. L. M. Bekkers, and E. M. G. van Esch. Immunologic factors involved in the malignant transformation of endometriosis to endometriosis-associated ovarian carcinoma. Cancer immunology, immunotherapy: CII, 70(7):1821–1829, July 2021.

24. Tiphaine Lefebvre, Manon Campas, Komodo Matta, Sadia Ouzia, Yann Guitton, Gauthier Duval, Stéphane Ploteau, Philippe Marchand, Bruno Le Bizec, Thomas Freour, Jean-Philippe Antignac, Pascal de Tullio, and German Cano-Sancho. A comprehensive multiplatform metabolomic analysis reveals alterations of 2-hydroxybutyric acid among women with deep endometriosis related to the pesticide trans-nonachlor. The Science of the Total Environment, 918:170678, March 2024.

25. Sebastian Lobentanzer, Patrick Aloy, Jan Baumbach, Balazs Bohar, Vincent J. Carey, Pornpimol Charoentong, Katharina Danhauser, Tunca Doğan, Johann Dreo, Ian Dunham, Elias Farr, Adrià Fernandez-Torras, Benjamin M. Gyori, Michael Hartung, Charles Tapley Hoyt, Christoph Klein, Tamas Korcsmaros, Andreas Maier, Matthias Mann, David Ochoa, Elena Pareja-Lorente, Ferdinand Popp, Martin Preusse, Niklas Probul, Benno Schwikowski, Bünyamin Sen, Maximilian T. Strauss, Denes Turei, Erva Ulusoy, Dagmar Waltemath, Judith A. H. Wodke, and Julio Saez-Rodriguez. Democratizing knowledge representation with BioCypher. Nature Biotechnology, 41(8):1056–1059, August 2023. Publisher: Nature Publishing Group.

26. Yuxing Lu, Sin Yee Goi, Xukai Zhao, and Jinzhuo Wang. Biomedical Knowledge Graph: A Survey of Domains, Tasks, and Real-World Applications, January 2025. 2501.11632 [cs].

27. Artem Lysenko, Irina A. Roznovăţ, Mansoor Saqi, Alexander Mazein, Christopher J. Rawlings, and Charles Auffray. Representing and querying disease networks using graph databases. BioData Mining, 9(1):23, July 2016.

28. Adnan Malik, Muhammad Arsalan, Carlos Moreno, Juan Mosquera, Eloy Félix, Tevfik Kizilören, Venkatesh Muthukrishnan, Barbara Zdrazil, Andrew R Leach, and Noel M O’Boyle. ChEBI: re-engineered for a sustainable future. Nucleic Acids Research, 54(D1):D1768–D1778, January 2026.

29. Jianping Man, Yufei Shi, Zhensheng Hu, Rui Yang, Zhisheng Huang, and Yi Zhou. KSDKG: construction and application of knowledge graph for kidney stone disease based on biomedical literature and public databases. Health Information Science and Systems, 12(1):54, December 2024.

30. Komodo Matta, Meriem Koual, Stéphane Ploteau, Xavier Coumoul, Karine Audouze, Bruno Le Bizec, Jean-Philippe Antignac, and German Cano-Sancho. Associations between Exposure to Organochlorine Chemicals and Endometriosis: A Systematic Review of Experimental Studies and Integration of Epidemiological Evidence. Environmental Health Perspectives, 129(7):76003, July 2021.

31. Komodo Matta, Tiphaine Lefebvre, Evelyne Vigneau, Véronique Cariou, Philippe Marchand, Yann Guitton, Anne-Lise Royer, Stéphane Ploteau, Bruno Le Bizec, Jean-Philippe Antignac, and German Cano-Sancho. Associations between persistent organic pollutants and endometriosis: A multiblock approach integrating metabolic and cytokine profiling. Environment International, 158:106926, January 2022.

32. Tilektes Maulenkul, Alina Kuandyk, Dinara Makhadiyeva, Anar Dautova, Milan Terzic, Ainash Oshibayeva, Ikilas Moldaliyev, Ardak Ayazbekov, Talgat Maimakov, Yerbolat Saruarov, Faye Foster, and Antonio Sarria-Santamera. Understanding the impact of endometriosis on women’s life: an integrative review of systematic reviews. BMC women’s health, 24(1):524, September 2024.

33. Dimitrios Michail, Joris Kinable, Barak Naveh, and John V. Sichi. Jgrapht—a java library for graph data structures and algorithms. ACM Trans. Math. Softw., 46(2), May 2020.

34. Alistair Miles and Sean Bechhofer. SKOS Simple Knowledge Organization System Reference. W3C Recommendation. World Wide Web Consortium, United States, August 2009.

35. Shawn T. O’Neil, Brian M. Schilder, Kevin Schaper, Corey Cox, Daniel Korn, Sarah Gehrke, Christopher J. Mungall, and Melissa A. Haendel. monarchr: An R Package for Querying Biomedical Knowledge Graphs. Bioinformatics (Oxford, England), page btaf549, September 2025.

36. Silvio Peroni and David Shotton. FaBiO and CiTO: Ontologies for describing bibliographic resources and citations. Journal of Web Semantics, 17:33–43, December 2012.

37. David Powers. Evaluation: From Precision, Recall and F-Factor to ROC, Informedness, Markedness & Correlation. Mach. Learn. Technol., 2, January 2008.

38. Kazuaki Suda, Hirofumi Nakaoka, Kosuke Yoshihara, Tatsuya Ishiguro, Ryo Tamura, Yutaro Mori, Kaoru Yamawaki, Sosuke Adachi, Tomoko Takahashi, Hiroaki Kase, Kenichi Tanaka, Tadashi Yamamoto, Teiichi Motoyama, Ituro Inoue, and Takayuki Enomoto. Clonal Expansion and Diversification of Cancer-Associated Mutations in Endometriosis and Normal Endometrium. Cell Reports, 24(7):1777–1789, August 2018.

39. DR Swanson. Medical literature as a potential source of new knowledge. Bulletin of the Medical Library Association, 78(1):29–37, January 1990.

40. Magdalena Szumilas. Explaining Odds Ratios. Journal of the Canadian Academy of Child and Adolescent Psychiatry, 19(3):227–229, August 2010.

41. D.R. Unni, S.A.T. Moxon, M. Bada, M. Brush, R. Bruskiewich, J.H. Caufield, P.A. Clemons, V. Dancik, M. Dumontier, K. Fecho, G. Glusman, J.J. Hadlock, N.L. Harris, A. Joshi, T. Putman, G.R. Qin, S.A. Ramsey, K.A. Shefchek, H. Solbrig, K. Soman, A.E. Thessen, M.A. Haendel, C. Bizon, C.J. Mungall, The Biomedical Data Translator Consortium, and Remzi Celebi. Biolink Model: A universal schema for knowledge graphs in clinical, biomedical, and translational science. Clinical and Translational Science, 15(8):1848–1855, August 2022.

42. Graham J. G. Upton. Fisher’s Exact Test. Journal of the Royal Statistical Society: Series A (Statistics in Society), 155(3):395–402, 1992. eprint: https://rss.onlinelibrary.wiley.com/doi/pdf/10.2307/2982890.

